# Transition between functional regimes in an integrate-and-fire network model of the thalamus

**DOI:** 10.1101/034918

**Authors:** Alessandro Barardi, Jordi Garcia-Ojalvo, Alberto Mazzoni

**Affiliations:** Departament of Experimental and Health Sciences, Universitat Pompeu Fabra, Dr. Aiguader 88, 08003 Barcelona, Spain; Departament de Física i Enginyeria Nuclear, Universitat Politècnica de Catalunya, Rambla Sant Nebridi 22, 08222 Terrassa, Spain; The BioRobotics Institute, Scuola Superiore Sant’Anna, Pontedera, 56026, Italy

## Abstract

The thalamus is a key brain element in the processing of sensory information. During the sleep and awake states, this brain area is characterized by the presence of two distinct dynamical regimes: in the sleep state activity is dominated by spindle oscillations (7 – 15 Hz) weakly affected by external stimuli, while in the awake state the activity is primarily driven by external stimuli. Here we develop a simple and computationally efficient model of the thalamus that exhibits two dynamical regimes with different information-processing capabilities, and study the transition between them. The network model includes glutamatergic thalamocortical (TC) relay neurons and gabaergic reticular (RE) neurons described by adaptative integrate-and-fire models in which spikes are induced by either depolarization or hyperpolarization rebound. We found a range of connectivity conditions under which the thalamic network composed by these neurons displays the two aforementioned dynamical regimes. Our results show that TC-RE loops generate spindle-like oscillations and that a critical value of clustering in the RE-RE connections is necessary for the coexistence of the two regimes. We also observe that the transition between the two regimes occurs when the external excitatory input on TC neurons (mimicking sensory stimulation) is large enough to cause a significant fraction of them to switch from hyperpolarization-rebound-driven firing to depolarization-driven firing. Overall, our model gives a novel and clear description of the role that the two types of neurons and their connectivity play in the dynamical regimes observed in the thalamus, and in the transition between them. These results pave the way for the development of efficient models of the transmission of sensory information from periphery to cortex.

**Author Summary:** The thalamus is known to exhibit two clearly distinct dynamical regimes with different functionalities. During slow-wave sleep the thalamus is dominated by internal activity and is hardly sensitive to external stimuli. In contrast, in the awake state, the thalamus modulates its activity according to the stimuli coming from the periphery. Here we study the conditions regulating the transition between these two states. To that end we implement a simple yet biologically realistic neuronal network model of the thalamus, based on single-neuron models that reproduce the properties of the two prominent types of thalamic neurons, namely thalamocortical relay cells and reticular neurons. We found that when reticular neurons are clustered the network exhibits two distinct dynamical regimes; one dominated by oscillations and insensitive to external stimuli (like sleep) and one sensitive to them (like wake). Moreover we found that the transition between the two regimes is due to the increase of the external excitatory input (corresponding to stronger sensory stimuli).

## Introduction

The thalamus is often identified as a relay station between subcortical and cortical areas, since all sensory pathways of the nervous system pass through it before reaching the cortex. Indeed, sensory inputs from visual, auditory and somato-sensory receptors reach the cortex through synapses on thalamocortical relay neurons in a specific region of the thalamus, which in turn projects into the corresponding area in the primary visual cortex. Along with these forward projections, there are local inhibitory neurons receiving inputs from feedback fibers from layer 6 to the corresponding thalamic nuclei [1]. It is thus reasonable to think that thalamus does not limit its activity to faithfully transmitting information to the cortex, but it might play a role in gating and modulating the flow of information towards the cortex [2–4], i.e. in selecting which external information is supposed to reach the cortex and when. In particular, this view is coherent with the important role found to be played by the thalamus in sleep/arousal/wake process [5–7], and attention [8–10].

The main kind of excitatory neurons in the thalamus are the above-mentioned thalamocortical relay (TC) neurons. In vitro studies [11, 12] have revealed that these neurons can operate in different firing modes depending on their voltage level. Near the resting membrane potential, TC neurons can produce trains of spikes with frequency proportional to the amplitude of the injected current, due to voltage-dependent currents that generate action potentials [1]. This is usually called *tonic mode*. Alternatively, when TC neurons are hyperpolarized they can operate in a *bursting mode*, characterized by high-frequency bursts of action potentials (300 Hz) in response to hyperpolarization.

During slow-wave sleep, TC neurons display strong spindle oscillations (7 – 15 Hz) independently from external stimuli [1, 13]. In contrast, in the awake state TC neurons are known to vary their activity according to inputs coming from the associated receptor layers, and to affect in turn the activity of the associated primary sensory cortex. For instance, TC neurons belonging to the lateral geniculate nucleus (LGN) and the ventral posterior nucleus (VPN) are modulated by the retina [14] and by the tactile afferents [15], respectively, and modulate in turn the activity of primary visual and somatosensory cortical areas [4, 16, 17]. TC neurons are also key components of the above-mentioned gating role of the thalamus, contributing to the selection of salient information during selective attention [10].

As suggested by Crick in his seminal paper [2], the role of modulating the efficacy of sensory transmission of TC neurons is mainly played by the neurons of the reticular nucleus of the thalamus (RE neurons). In particular, the activation of RE neurons can strongly hyperpolarize TC neurons, which consequently undergo inhibitory rebound that gives rise to an endogenous oscillatory activity [18]. Specifically, spindles can be originated by TC bursts eliciting firing activity in RE cells. In turn RE bursts hyperpolarize TC cells, which consequently stop firing. When RE cells, lacking excitatory drive, stop firing too, the rebound of TC cells from hyperpolarization causes them to emit a burst of spikes and the cycle starts again. The overall process takes about 100 ms and generates rhythmic spindle oscillations. Therefore spindle generation is due to an interplay between TC and RE cells [19, 20]. Coherently with this fact, manipulating the activity of RE neurons was found to have behavioral consequences in attention tasks [10, 21].

During the awake state, TC cells undergo a transition and alternate this bursting mode with a tonic mode. As mentioned above, both modes are typical of TC neurons, and they could provide different frameworks for information processing, since during the bursting mode action potentials in the TC cell are not linked directly to EPSPs in that cell, whereas the opposite is true in the tonic mode. Therefore we expect that the bursting mode transmits information less efficiently than the tonic mode, in which an increase in the extra-thalamic inputs on TC neurons leads to a direct increase in the response of TC neurons [3]. How the thalamus exhibits the functional transition between the two regimes is not clear. In fact, a coherent view accounting for both TC and RE interactions and the resulting functional behavior of the thalamic network is still missing, due in particular to the relative paucity of simultaneous neurophysiological recordings of the two neuron types *in vivo*. In this context, the role of modeling becomes very relevant, due to its capacity to suggest candidate mechanisms for the generation of the observed behavior. Modeling of thalamic networks has been investigated for more than 20 years [22, 23], during which network models have been developed that capture a wealth of thalamic phenomena [24]. However, almost all studies to date have adopted neuron models at least as complex as the Hodgkin and Huxley model [22, 25], probably due to the aforementioned role of rebound currents. We are aware of only one attempt to model realistically thalamic interactions with integrate-and-fire (IF) neurons [26].

Here, we decided to focus on a single property of the thalamic network as a whole, namely its above-mentioned ability to switch between two dynamical regimes that display different external input sensitivity. We also study the role played in this phenomenon by the network architecture (connectivity and synaptic strength), ranging from loops of two neurons to the effect of sensory and cortical input on the whole thalamic network. To that end we have developed a thalamus TC-RE network model based on a particularly simple spiking neuron model, namely a different suited version of the adaptive exponential integrate-and-fire (aeIF) neuron model [27] for each neuron type. In the Results section we build our network progressively. First, we show how our aeIF neurons reproduce the two activity modes of TC and RE neurons: the standard depolarizing regime [12] and the rebound from hyperpolarization [18]. Then we investigate how spindle oscillations are generated through TC-RE interaction as a function of their coupling and of the presence of external inputs, and how heterogeneity can be tamed by the interaction of different TC-RE loops. Analysis of the complete network leads to our two main results: (i) a critical value of RE-RE clustering favors the presence of large scale spindle oscillations, and (ii) in the presence of clustering the network displays two dynamical regimes as the sensory input increases: it is insensitive to stimuli below a given intensity threshold, while above this threshold TC neurons (but not RE neurons) modulate their activity as a function of the input. Finally, we test that these conclusions hold also in the presence of cortical inputs impinging on the reticular neurons.

## Results

The presence of two different dynamical regimes in the thalamus has been known for decades [28–30]. This behaviour can be linked to a specific property of the two main kinds of neurons in the thalamus, described above, glutamatergic thalamocortical relay (TC) neurons and gabaergic thalamic reticular (RE) neurons. Both types of neurons can fire either as a result of depolarizing driving or as a rebound due to hyperpolarizing driving. In the following we will show how we modified an existing (aeIF) model of thalamic neurons [26] to reproduce the two types of responses for both kinds of neurons (Fig. 1). We investigated how and when the connectivity between the neurons displaying these properties induces a regime dominated by spindle oscillations, or responding to stimuli in a tonic-like mode. The analysis starts from two-neuron loops and extends up to full networks receiving input from the periphery and the cortical areas.

**Figure 1.**
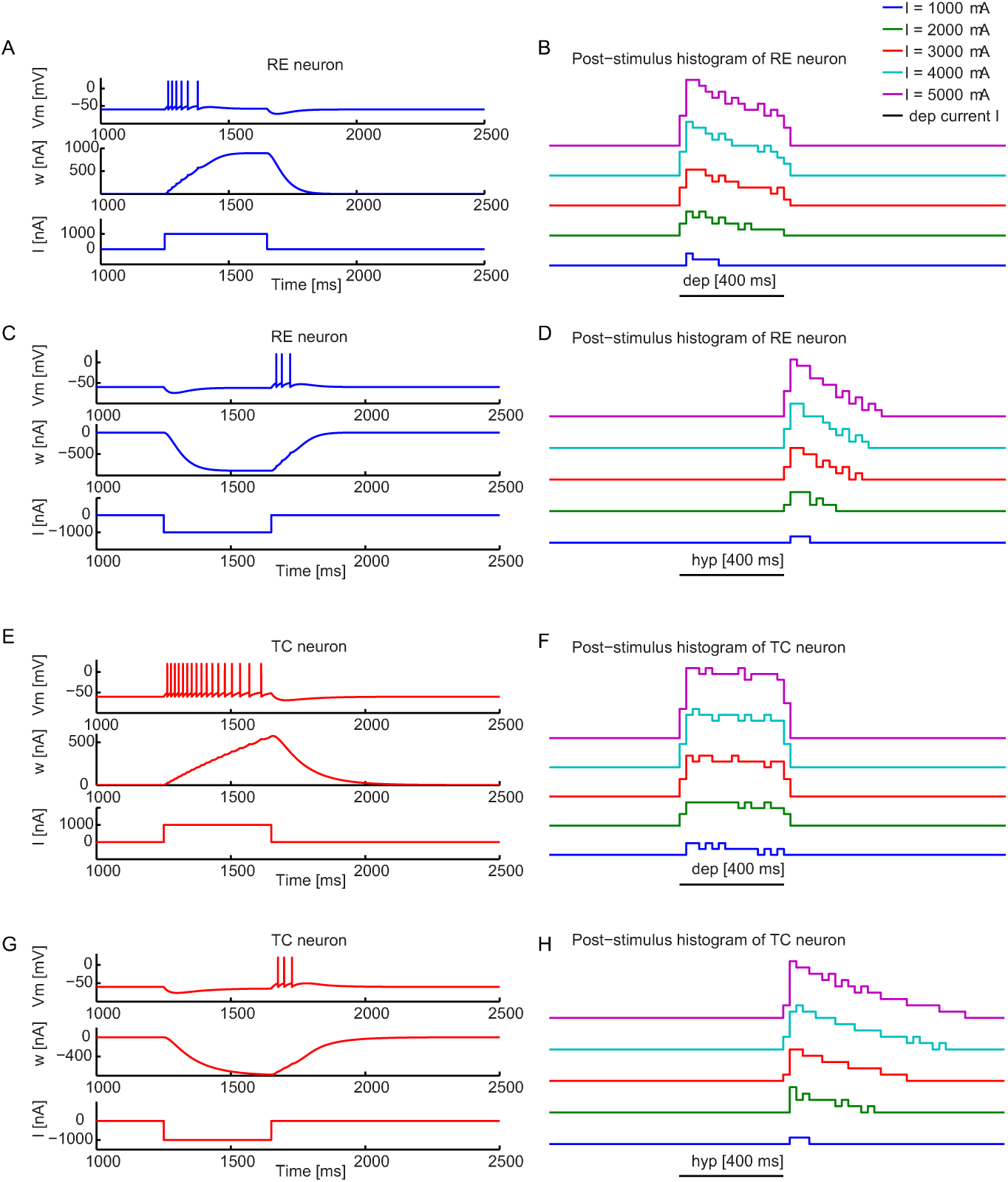
Dynamical properties of single RE and TC neurons as a function of input current. (A) Depolarization activity of a RE neuron. Membrane voltage (top) and adaptation variable (middle) of a RE neuron in response to a depolarizing current (bottom). (B) Corresponding post-stimulus time histograms for increasing depolarizing currents. (C) Hyperpolarization-rebound activity of a RE neuron and (D) corresponding post-stimulus time histograms for increasing hyperpolarizing currents. Parameters *a* and *b*, representing respectively the dynamics and the strength of adaptation (see equation 2) of RE neurons are defined in this way: *a* = 0.4 *μ*S and *b* = 0.02 nA. (E) Depolarization activity of a TC neuron and (F) corresponding post-stimulus time histograms for increasing depolarizing currents. (G) Hyperpolarization-rebound activity of a TC neuron and (H) corresponding post-stimulus time histograms for increasing hyperpolarizing currents. The values *a* and *b* are 0.2 *μ*S and 0 nA. The current intensity in (A, C, E, G) is 1000 mA, while it varies between 1000 mA and 5000 mA in panels (B, D, F, H). *V_T_* = −50 mV is the threshold potential for both types of neurons. Other parameters are defined in the Materials and Methods section.

### Dynamics of single neurons

The first step towards reproducing the two dynamical regimes of the thalamus described above, and the transition between them, is to choose a single-neuron model able to capture the peculiar properties of thalamic neurons, and in particular the firing induced by hyperpolarization-driven rebound. To that end we selected a properly tuned adaptative exponential integrate-and-fire (aeIF) spiking neuron model [27, 31, 32] (see Materials and Methods Section) for each of the two thalamic neuron types considered. By tuning the key parameters of the aeIF model it is possible to adjust the dynamics and the strength of adaptation (parameters *a* and *b* in Eq. 2, respectively, in the Materials and Methods Section) to reproduce the intrinsic dynamical modes typical of thalamic neurons.

For *a* = 0.4 *μ*S and *b* = 0.02 nA, the RE aeIF neuron models (RE neuron from now on) exhibits regular firing activity in response to depolarizing stimuli (Fig. 1A, B), while they display bursting activity in response to hyperpolarizing stimuli (Fig. 1C, D), consistently with experimental findings [33–35]. In particular, in response to a depolarizing stimulus (Fig. 1A), RE neurons display firing activity with a certain degree of spike-frequency adaptation that saturates before the end of the stimulus and stops neuronal firing. For large enough applied currents, the response extends for the whole duration of the stimulus (Fig. 1B). In response to a hyperpolarizing stimulus (Fig. 1C, D), and due to the relatively large value of *a*, the neuron exhibits rebound bursting activity, also with spike-frequency adaptation, for the same spike threshold used in the depolarizing case.

TC neurons generally show a more robust bursting activity and a negligible level of spike-frequency adaptation [11] (see [1] for a review). This is achieved in the model by imposing a larger value of *a* = 0.2 *μ*S and *b* = 0 nA, thus making the adaptation strength negligible. In particular, in response to a depolarizing stimulus our TC neurons model produce patterns of firing activity (Fig. 1E) with negligible spike-frequency adaptation (Fig. 1F) (leading thereby to high firing activity for all the duration of the stimulus). In contrast, a hyperpolarizing stimulus leads to rebound bursting (Fig. 1G) and moderate spike-frequency adaptation (larger than in RE neurons) (Fig. 1H). In the case of depolarizing stimuli, characterized by negligible adaptation and regular firing activity, TC neurons exhibit an effective increase of activity (Fig. 1F) according to the increasing external input and compatibly with the refractory period, where neuron is not allowed to fire. Therefore the firing activity increases proportionally with larger external sensory inputs. This is consistent with the linear input-output relation in the tonic mode (Fig. 1E, F), in contrast with the bursting mode where there is no direct link between the EPSP and spike generation, which thus corresponds to a nonlinear input-output relation [36].

Overall these results show that the aeIF models properly capture the two firing modes (depolarizing-driven and hyperpolarization-driven) for both TC and RE neurons. In the following we will show the transition between the two modes for TC neurons due to external inputs, and how recurrent activity drives a transition at the network level from stimulus-insensitive to stimulus-sensitive behavior.

### Two-neuron loops

Before moving to large, structured networks we carefully analyzed the properties of the mutual interaction between TC and RE neurons. Specifically, we studied different simple two-neuron loops formed by TC-RE and RE-RE neurons, and examined how self-sustained oscillatory patterns originated in these networks are modulated by synaptic strengths regulating the internal recurrent activity. We also studied the effect of GABA temporal decay dynamics on the frequency of oscillation, and the input-driven oscillatory pattern of a TC-RE loop. This analysis is informative towards the building of a full network.

We first built a minimal model of two bidirectionally coupled neurons, a RE neuron and a TC neuron (Fig. 2A). Activating this RE-TC loop for 50 ms leads to oscillations that persist stably after the stimulus termination (Fig. 2B). These oscillations are due to the rebound bursting properties of the TC relay cell, which is mutually connected with the RE neuron: the TC neuron provides depolarizing input to the RE neuron, which displays bursting activity that generates strong hyperpolarization, followed by rebound firing activity in TC neurons. Consequently, in this configuration the RE neuron fires in response to depolarizing currents, while the TC neuron fires only in response to hyperpolarizing inputs.

**Figure 2.**
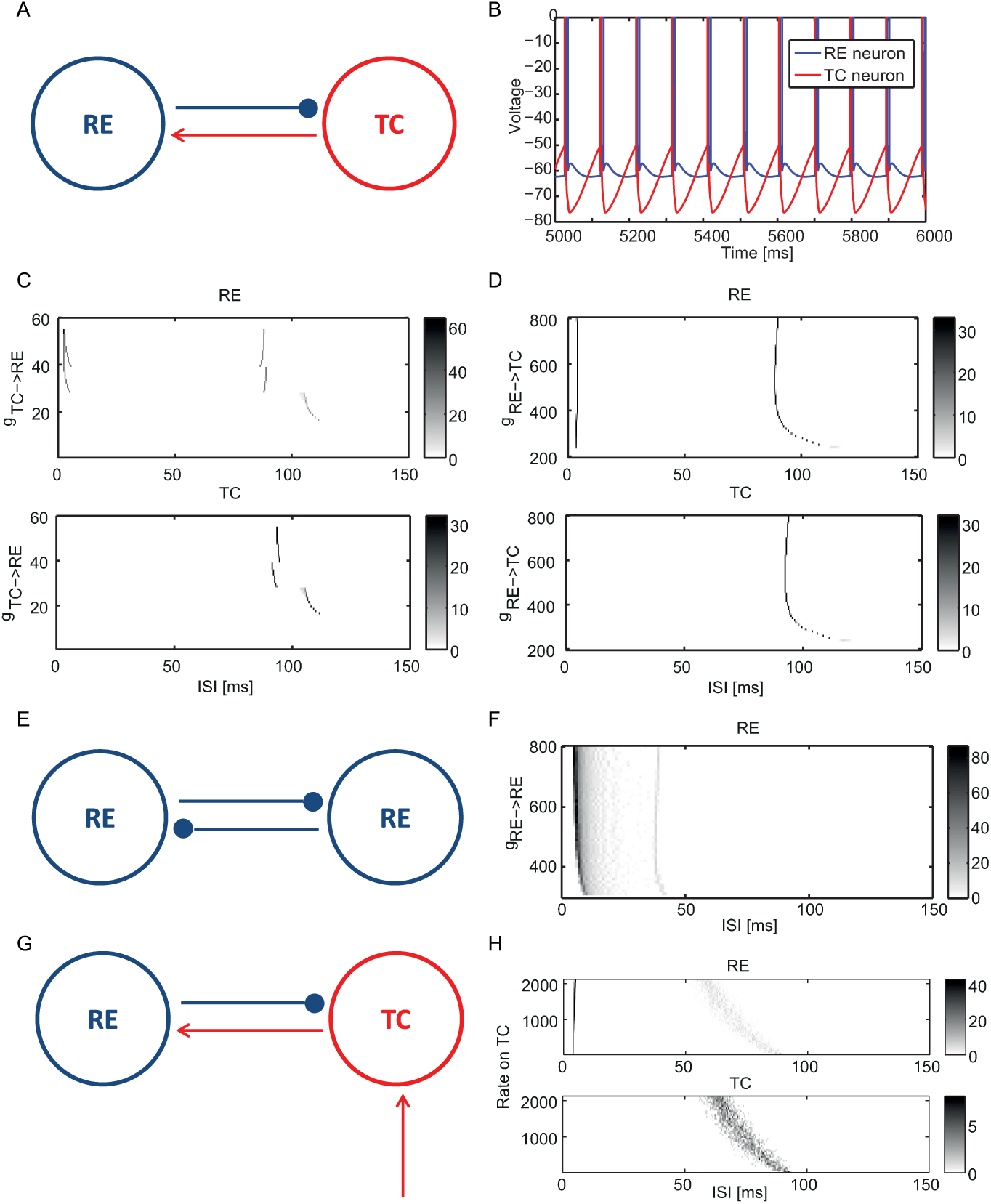
Dynamical properties of two-neuron loops. (A) Scheme of a two-neuron TC-RE loop. (B) Membrane voltage traces of the TC and RE neurons generated by this minimal TC-RE loop. (C) Interspike interval (ISI) distribution of the TC-RE loop as a function of the synaptic strength *g_TC→RE_*. The value of *g_RE_*_→_*_TC_* is appropriately set to 550 *μ*S in order to support self-sustained activity, while *g_TC→RE_* varies between 10 *μ*S and 60 *μ*S. RE and TC ISI distributions are shown in the top and bottom plots, respectively. (D) ISI distribution of a TC-RE loop as a function of the synaptic strength *g_RE_*_→_*_TC_*. The value of *g_TC→RE_* is chosen equal to 32 *μ*S to reproduce the two-spike bursting dynamical regime of panel *b* while *g_RE_*_→_*_TC_* varies between 200 and 800 *μ*S. RE and TC ISI distributions are shown in the top and bottom plots, respectively. (E) Scheme of a minimal purely reticular RE-RE loop. (F) ISI distribution of this loop as a function of the synaptic strength *g_RE_*_→_*_RE_*. *g_RE_*_→_*_RE_* varies between 200 *μ*S and 800 *μ*S. (G) Scheme of an input-driven two-neuron TC-RE loop. (H) ISI distribution of this loop as a function of external sensory input strength. RE and TC ISI distributions are shown in the top and bottom plots, respectively. The synaptic strengths are respectively: *g_RE_*_→_*_TC_* = 550 *μ*S, *g_TC→RE_* = 32 *μ*S and *g_EXT→TC_* = 1 *μ*S.

Next we investigated how these oscillatory patterns vary as a function of the synaptic strength *g_TC→RE_*, keeping *g_RE_*_→_*_TC_* to a reference value of 550 *μ*S. By increasing *g_TC→RE_*, both the TC and RE neurons oscillate with higher frequencies, as can be seen from the decrease of the inter-spike interval (ISI) in Fig. 2C (bottom). Stronger synaptic strengths enhance the firing activity of the RE neuron, which fires in advance along the oscillation cycle and thus leads the TC neuron to spike at an earlier phase. The net effect is an increase in the oscillation frequency. The RE neuron (Fig. 2C, top) displays bursting activity in response to depolarizing input above a threshold value of *g_TC→RE_* = 29 *μ*S. It oscillates at around 11 Hz (inter-burst ISI ≈ 90 ms) with two spikes per burst with an intra-burst ISI ≈ 5 ms. By increasing the synaptic strength *g_TC→RE_*, the neuron passes a second threshold *g_TC→RE_* = 40 *μ*S and presents three spikes per burst (three ISIs are present), eventually entering a regime in which the ISI approaches the intrinsic refractory period of the neuron (2.5 ms, see Material and Methods section).

Subsequently we performed the complementary analysis by fixing *g_TC→RE_* to 32 *μ*S (which led to two-spike bursting in the preceding analysis) and varying *g_RE_*_→_*_TC_*. Figure 2D shows that as *g_RE_*_→_*_TC_* is increased, the TC neuron oscillates with a gradually increasing frequency that stabilizes around 10.5 Hz (Fig. 2D, bottom), while the RE neuron displays bursting activity with the same inter-burst ISI as the TC neuron and an intra-burst ISI of ≈ 3 ms (two-spikes-per-second scenario of previous analysis) (Fig. 2D. top). Note that the brief hyperpolarization induced in the TC cell by the firing of a single RE cell is able to trigger only one rebound spike, and consequently the number of spikes/burst in the RE cell remains constant. This is consistent with the results reported in Ref. [1], where spindle activity required at least a four-neuron network (see next Section).

Next we explored the dynamics of a purely GABAergic reticular RE-RE loop (Fig. 2E) as a function of the synaptic strength *g_RE_*_→_*_RE_*. As Fig. 2F shows, the RE neurons present a sustained strong and adapting bursting activity (corresponding to a wide range of intraburst ISI) and for increasing values of the synaptic strength, the inter-burst ISI decreases. Importantly, unlike in previous studies, here the decreasing inter-burst ISI does not entail an increase in oscillation frequency, since here bursts last much longer (with more than 10 spikes per burst). This result shows that RE-RE synapses strengthen the rebound bursting properties and can be expected to enhance the bursting activity in a larger network.

In the simple TC-RE loop motif, the oscillation frequency can be tuned by the GABA decay time constant. For instance, by varying *τ_decay_* from 5 to 35 ms in the minimal model of Fig. 2D, the frequency of the two neurons oscillates between ~ 25 and 6 Hz (Supp. Fig. S1). This leads to corresponding changes in the ISI distributions (Figs. S2–S3), without qualitative variations with respect to the behavior shown in Fig. 2.

After investigating the properties of stand-alone RE-TC loops, we moved to analyze an input-driven loop in which the TC neuron receives an external sensory input modeled as a Poisson distribution with increasing amplitude (Fig. 2G). We only considered inputs to TC, mimicking the sensory stimuli coming from the retina or the peripheral nervous system. We set reference values of *g_TC→RE_* = 32 *μ*S and *g_RE_*_→_*_TC_* = 550 *μ*S and GABA *τ_decay_* = 20ms, for which the spontaneous activity (in the absence of external input) corresponds to low-frequency bursting with two spikes per burst. The value of GABA *τ_decay_* is lower in the full population model. When we increased the external input rate (Fig. 2H) the ISI distribution was significantly different from the one observed in the absence of external stimulus (Fig. 2D): both neurons show a strong variation in the bursting frequency due to the external stimulus, and the ISI displays a large variance due to the introduction of noise. On the other hand, and consistently with Fig. 2D, the RE neuron is in bursting mode for all values of external input, with the ISI approaching the refractory period.

### Four-neuron motifs

As a last step before moving to the full network, we investigated several four-neuron motifs, made of two RE and two TC neurons, to understand what are the structural connectivity features more suitable to explain large oscillatory synchronization phenomena, namely spindle oscillations, in the bursting regime, even in presence of heterogeneity between neurons. Previous work has shown [26] that aeIF models are able to reproduce this self-sustained oscillatory behavior in the form of periodic bursting, and that the minimal circuit reproducing phenomen is a circuit of two TC and two RE neurons fully connected with each other, with the exception of TC-TC connections, which are not present in the thalamus [37]. As in the case of the two-neuron loop, bursting is mainly due to the rebound bursting properties of TC cells and RE cells (Fig. 3F) [1], and the oscillation frequency depends on the GABA temporal decay constant.

**Figure 3.**
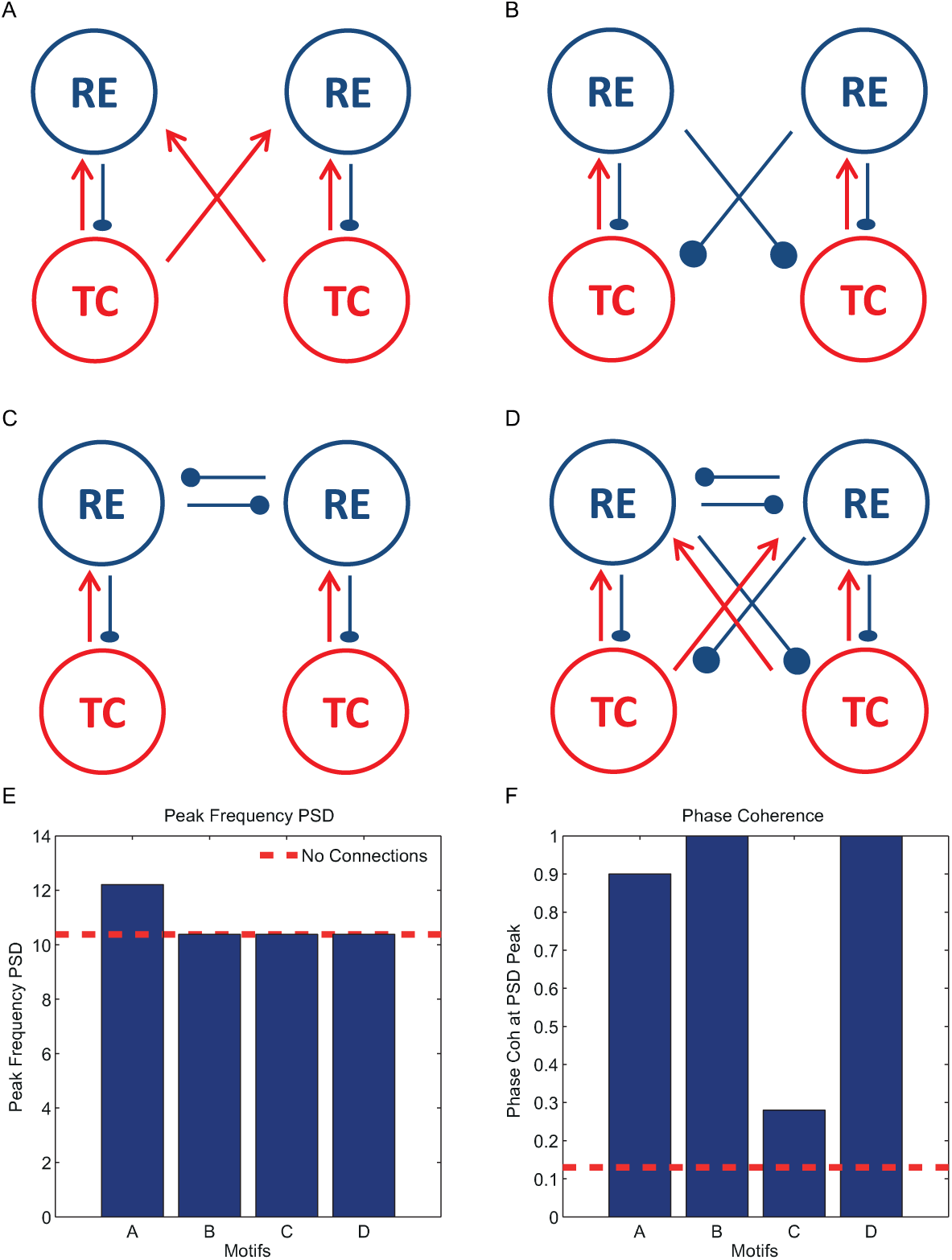
Four-neuron motifs in the form of coupled pairs of TC-RE loops. The two TC-RE oscillators are bidirectionally coupled through (A) TC-RE connections, (B) RE-TC connections, (C) RE-RE connections, and (D) all three connections. (E) Frequency of the power spectral peak and (F) phase coherence at that frequency for the four different motifs. The power spectral density and phase coherence were averaged across 50 trials for random values of the GABA decay time (see text). GABA rise time and AMPA rise and decay times are set constant (see Materials and Methods section). When the corresponding connections exist in the motifs, the synaptic strengths are respectively: *g_RE→TC_* = 550 *μ*S, *g_TC→RE_* = 32 *μ*S and *g_RE→RE_* = 20 *μ*S.

We studied different couplings between pairs of two-neuron TC-RE loops (which are equivalent to two bidirectionally coupled oscillators), and analyzed which coupling configuration leads more readily to oscillatory spindle patterns by examining the power spectrum of TC neurons and the phase coherence between them. Figure 3 shows the schemes of the different circuits explored depending on the coupling links being considered: TC-RE connections (Fig. 3A), RE-TC connections (Fig. 3B), RE-RE connections (Fig. 3C) and all three types of connections (Fig. 3D). For each circuit, we calculated the power spectral density and phase coherence between the two loops (see Materials and Methods section) by using the activity of TC neurons. The phase coherence is calculated by averaging 50 trials each with a different GABA *τ_decay_* drawn from a Gaussian distribution with mean 20 ms and standard deviation 5 ms, which leads to variability in the frequencies of the two TC-RE loops being coupled.

Figure 3E shows the frequency at which the power spectrum of the TC neuron activity has its maximum, and Fig. 3F the corresponding phase coherence at that frequency. The horizontal dashed red lines represent the corresponding values in the case of uncoupled loops. In the uncoupled case, the oscillation frequency is ≈ 10.4Hz and the loops are weakly synchronized (the phase coherence being ≈ 0.12). The two TC-RE oscillators strongly synchronize with a zero-lag phase (corresponding time lag is ≈ 0, not shown) with respect to the uncoupled case, while the loops are poorly zero-lag synchronized when only RE-RE connections are present. Therefore this result supports the idea that spindle generation is mainly due to an interplay between TC and RE cells [19, 20], which is enhanced by RE-RE connections.

### Full thalamic network

We finally extended the size of the network to 500 neurons to capture the dynamics of a complex thalamic structure. Following experimental indications [38–40], we consider that each RE projects four connections to TC neurons and to RE neurons themselves, while TC neurons have only on average one connection with RE neurons only. The GABA decay time is set to *τ_gaba_* = 10 ms. The structural connectivity is built according to the *small world algorithm* of Strogatz and Watts [41], to reproduce the fact that brain circuits have local modularity and long-range connectivity [42]. We first investigated to what extent the oscillations in the spindle frequency range (7–15 Hz) in both TC and RE neurons are robust to the introduction of a certain degree of inter-clustering between RE-RE neurons. Figure 4A shows the connectivity matrix of a random network with rewiring probability *RP* = 1, and Fig. 4B that of a network with rewiring probability *RP* = 0.25.

**Figure 4.**
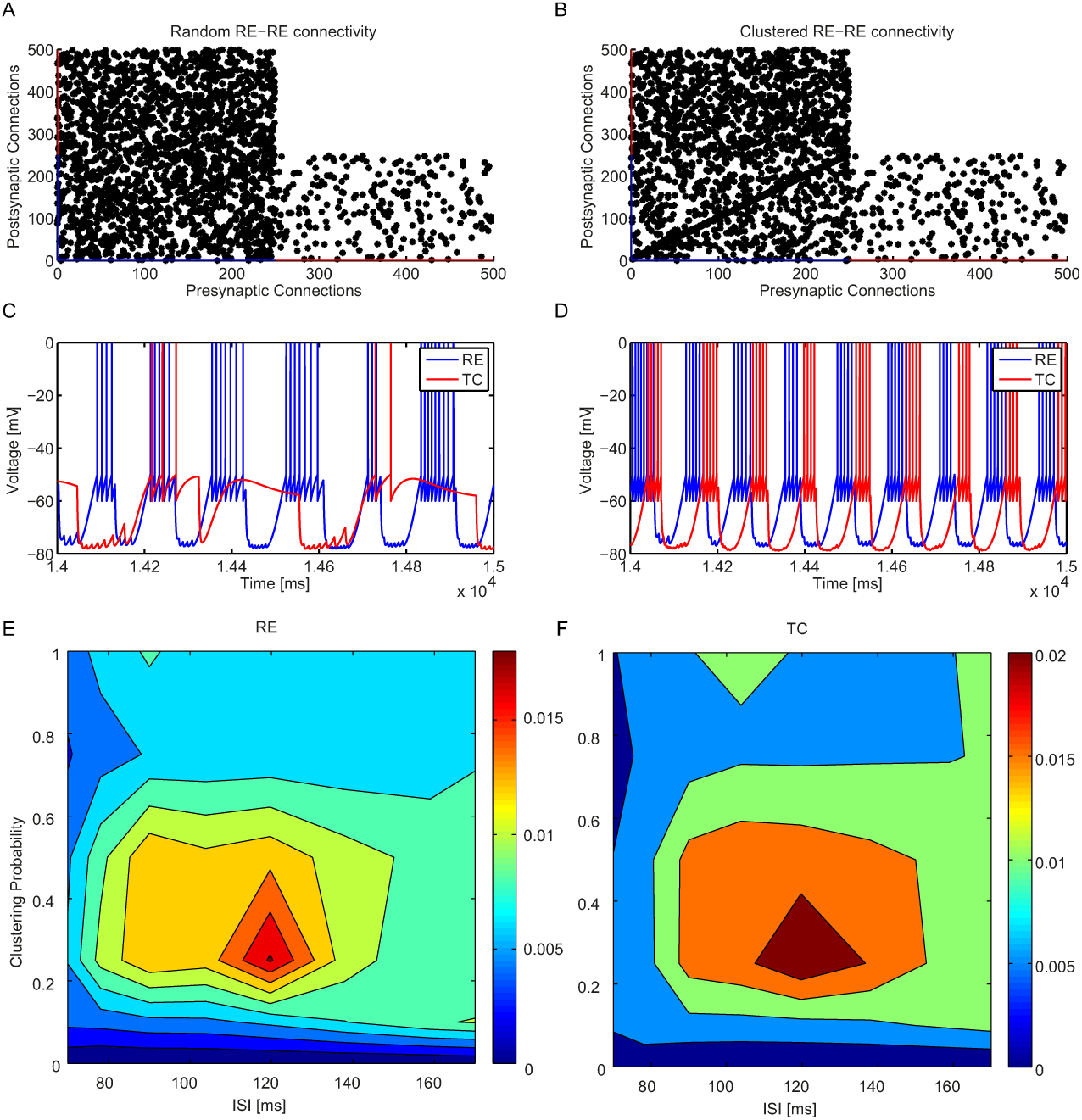
Spindle activity generated by a full network of TC-RE neurons depending on RE-RE clustering. (A) Connectivity matrix of a random TC-RE network. The presynaptic neurons are represented in the x axis and the postsynaptic neurons in the y axis. The network is made of 500 neurons, of which the first 250 are RE neurons and the remaining ones are TC neurons. (B) Connectivity matrix in the presence of RE-RE clustering (rewiring probability *RP* = 0.25) (C) Membrane voltage dynamics of a couple of arbitrarily chosen TC and RE neurons in the case of random network. (D) Membrane voltage dynamics of a couple of arbitrarily chosen TC and RE neurons in the presence of clustering: evidence of typical spindle oscillations. (E) ISI distribution (color-coded) as a function of the rewiring probability for RE (left) and TC (right) neurons. The synaptic strengths are respectively: *g_RE_*_→_*_TC_* = 300 *μ*S, *g_TC→RE_* = 200 *μ*S and *g_RE→RE_* = 300 *μ*S.

We found that in the random network (Fig. 4A), temporally irregular bursting is dominant (Fig. 4C). On the other hand, in the presence of RE-RE clustering (Fig. 4B) the network shows quite regular and synchronized spindle oscillations at 8 Hz (Figure 4D). In order to characterize and quantify the bursting regular state (or spindle rhythm) and distinguish it from irregular tonic activity, we studied the inter-burst interval distribution (in particular the probability of a peak of ISI distribution above 50 ms) as a function of the rewiring probability *RP* of the small world architecture (see Methods for details). Our results, shown in Fig. 4E, reveal that fully regular networks (*RP* = 0, each neuron projects regularly to a fixed number of adjacent neurons) cannot support regular bursting activity and are often almost silent (with a firing rate of around 0.4 spikes/s, results not shown). At the other extreme, fully random networks (*RP* = 1) show sustained activity with temporally irregular bursting of TC and RE neurons. Between these two conditions, there is an optimum rewiring probability (*RP* ~ 0.25) showing a relatively large ISI peak corresponding to frequency ~ 8.5 Hz. The fraction of neurons displaying a large inter-burst ISI peak decreases substantially for increasing rewiring probability, namely when going towards fully random networks. Intuitively, given that connections between thalamic circuits are local but sparse [38–40], excitatory synapses are very sparse and they are more effective when they impinge on small clusters of RE-RE neurons, enhancing and modulating the oscillatory spindle rhythm.

Given the results obtained above, we decided to study a network with the critical degree of clustering (*RP* = 0.25), and simulate constant external sensory input of different intensities impinging on TC neurons. We tested if by increasing the external input on these neurons the network showed a transition from bursting to tonic mode, which could be associated with the switch from sleep to awake state [28–30]. Given the nonlinear relation between input and output in the bursting mode [36], we expect to see a change in the firing rate trend of TC neurons (the neurons that project to the cortex) only when the network goes from bursting to tonic, through which the firing rate should increase with the input. Figure 5A shows the firing rate of TC (red) and RE (blue) neurons for increasing external sensory input on TC neurons. The case of input *S* = 0 spikes/s corresponds to the self-sustained condition discussed above. By increasing the input amplitude, the network displays a transition in the firing rate of TC neurons at around *S* = 50 spikes/s, after which the response of the thalamus increases sub-linearly with the external input. We interpret this as an indication of the switch from a purely bursting mode to a temporally irregular state. Note that the driver of this transition is the response of the recurrent activity to the external sensory input, since we did not change the intrinsic parameters of the model.

**Figure 5.**
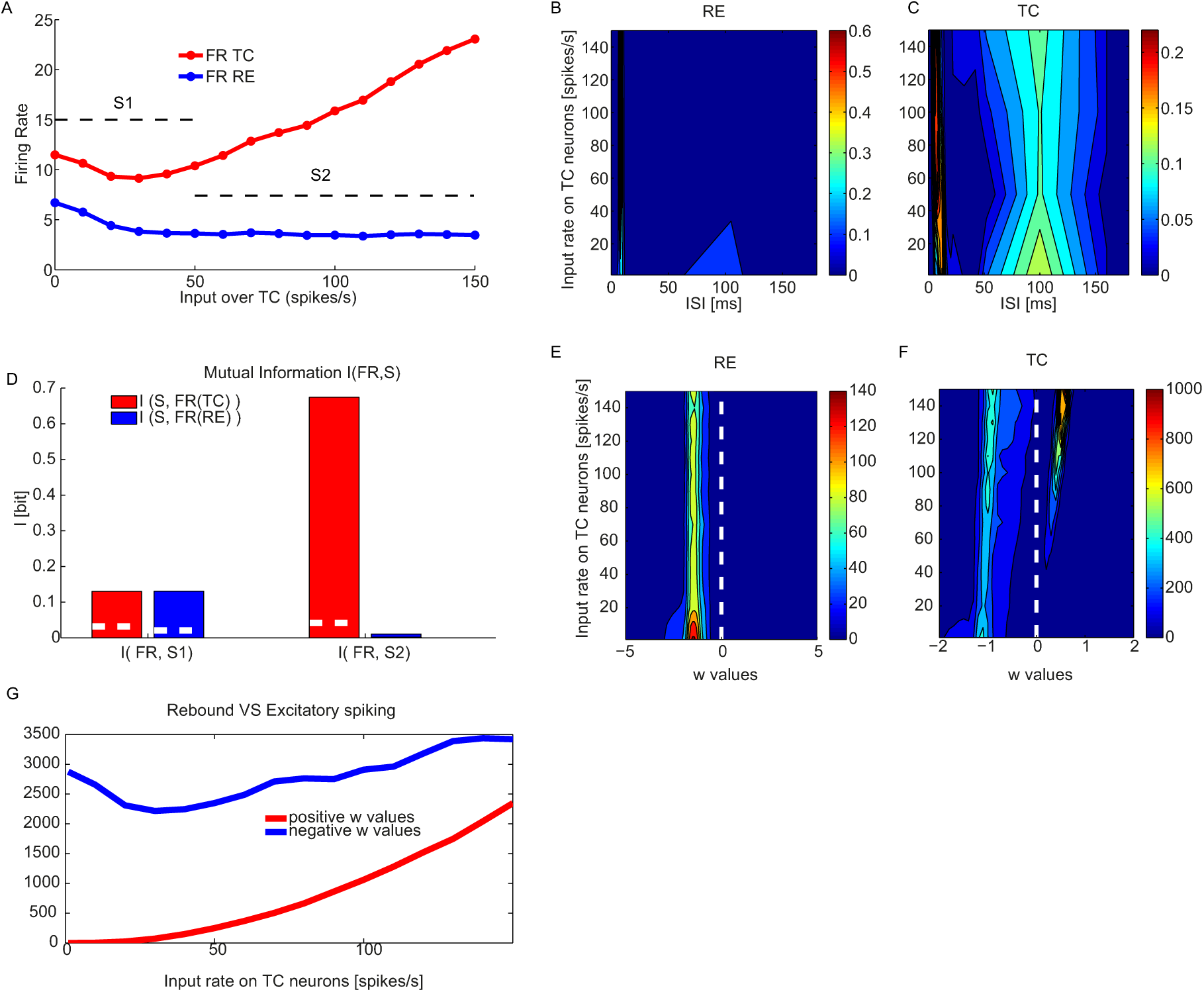
Bursting and tonic modes displayed by a TC-RE network with RE-RE clustering as a function of external input on TC neurons. (A) Firing rate of TC (red) and RE (blue) neurons as a function of external driving input impinging on TC neurons. (B, C) ISI distribution as a function of external driving input on TC neurons of RE (B) and TC (C) neurons. (D) Mutual Information between the set of increasing external stimulus (0–150 spikes/s) and the neural response given by the firing rate of TC and RE neurons. Different external sensory inputs are considered for the two regimes, following panel A: 0–50 spikes/s for the bursting mode and 60–150 spikes/s for the tonic mode. The white dashed line in the bar plots refers to significance threshold (*p <* 0.05, bootstrap test). The measures are averaged over 100 trials for each external stimulus. (E, F) Adaptation variable *w* of RE (E) and TC (F) neurons (color coded) as a function of the external input on TC neurons, averaged across 100 trials for each external stimulus. (G) Number of positive *w* values (depolarizing events) and negative *w* values (rebound events) of TC neurons. The synaptic strengths are respectively: *g_RE_*_→_*_TC_* = 300 *μ*S, *g_TC→RE_* = 200 *μ*S and *g_RE_*_→_*_RE_* = 300 *μ*S.

In order to explore this scenario further, we calculated the ISI distribution of RE and TC neurons by averaging over 100 trials for each different stimulus *S*. The RE neurons are the most insensitive to increasing external input, as can be seen in Fig. 5B. On the other hand the fraction of TC neurons displaying a large inter-burst ISI decreased as the stimulus intensity surpasses a critical value (going from region S1 to region S2 in Fig. 5A), and a corresponding increase of the intra-burst ISI peak approaching the refractory period (2.5 ms, see Materials and Methods section). We classified this as a further signature of a transition between a bursting mode and an irregular firing regime.

Next we calculated the information about the stimuli carried by the firing rates of the TC and RE neurons in the two different regimes. To that end we used the mutual information (see Methods), which quantifies the reduction of the uncertainty in predicting the applied stimulus given a single observation of the triggered response. In this case we considered a rate code, i.e. we selected as response the average firing rate over the whole stimulation [43]. Figure 5C compares *I*(*S*1; FR) and *I*(*S*2; *FR*) between the firing rates of TC (red) and RE (blue) neurons and the set of stimuli *S*1 and *S*2, where *S*1 ranges between 0 and 50 spikes/s, while *S*2 varies from 60 to 150 spikes/s, corresponding to the two dynamical regimes of Fig. 5A. The figure clearly shows that in the bursting mode both the RE and TC neurons carry a lower information (0.13 bit, *p <* 0.05: bootstrap test), in comparison with the information encoded by TC neurons during the tonic mode (≈ 0.7 bit, *p <* 0.05: bootstrap test). RE neurons during the tonic mode do not encode significant information, in fact their firing rate decreases with respect to the bursting regime and after that remains constant for all inputs. These results show that the information about the stimulus that the thalamus carries (and is then potentially able to convey to the cortex) is much higher in the tonic mode, since in that regime spontaneous activity is enhanced and this contributes to keeping an almost linear relation between input and output and thus to minimizing rectification of the response [36].

In order to further interpret this transition, we examined the nature of each TC and RE spike by checking the sign of the adaptation variable *w* at the spiking time of each neuron. A positive value of *w* indicates that neuron fires via a depolarizing input (see Fig. 1), while if negative we classify it is as a rebound spike. Fig. 5E shows that RE neurons spike mostly due to a rebound in response to hyperpolarizing inputs (coming only from internal RE-RE clustered connections) for all the range of sensory input over TC neurons. TC neurons, in turn, also fire mainly in response to incoming hyperpolarizing currents (in this case coming from RE neurons) during the burst mode (Fig. 5F), and after the transition from bursting to tonic mode a fraction of the spikes occur in response to depolarizing external inputs. Thus the transition occurring at around *S* = 50 spikes/s, shown in figure 5A, underlies a shift in the spiking mechanism profile. This is confirmed in Fig. 5G, which shows a quantitative estimation of the effective number of excitatory-driven spikes (blue) and inhibitory-rebound spikes (red) as the external input increases.

So far we have considered a thalamic network receiving an external sensory input impinging on TC neurons. We complete the picture including also a corticothalamic input [44] projecting to RE neurons. Figure 6A shows that the transition dynamics is not altered by the addition of a constant input from the cortex, which results only on an increase of the firing rate for both kind of neurons. The appearance and the increase of depolarization spikes occur for similar levels of inputs (Fig. 6B). The amount of information carried by RE and TC neurons in the two different regimes is relatively unaltered (Fig. 6C, D), supporting the hypothesis that the information carried by projecting neurons during the tonic mode is higher than in the bursting mode. Interestingly, by increasing the amplitude of the cortical input on RE (from 1000 to 2000 spikes/s), the information encoded by TC neurons is increased for the tonic mode (from 0.6 to 0.66 bit, *p <* 0.05, bootstrap test) (Fig. 6D). This result highlights the role of the intrinsic rebound bursting properties of TC neurons, which are essential in the generation of the spindle rhythm. They could also reinforce the role of corticothalamic feedback in information processing, for instance by recruiting TC neurons through inhibition and thus modulating TC firing rate [44]. To support the importance of rebound bursting properties of TC neurons, we plotted in Supp. Fig. S4 the firing rate of TC and RE neurons, the ISI distribution and the w distribution at a fixed rate of external sensory input on TC neurons (150 spikes/s), for different levels of cortical input.

**Figure 6.**
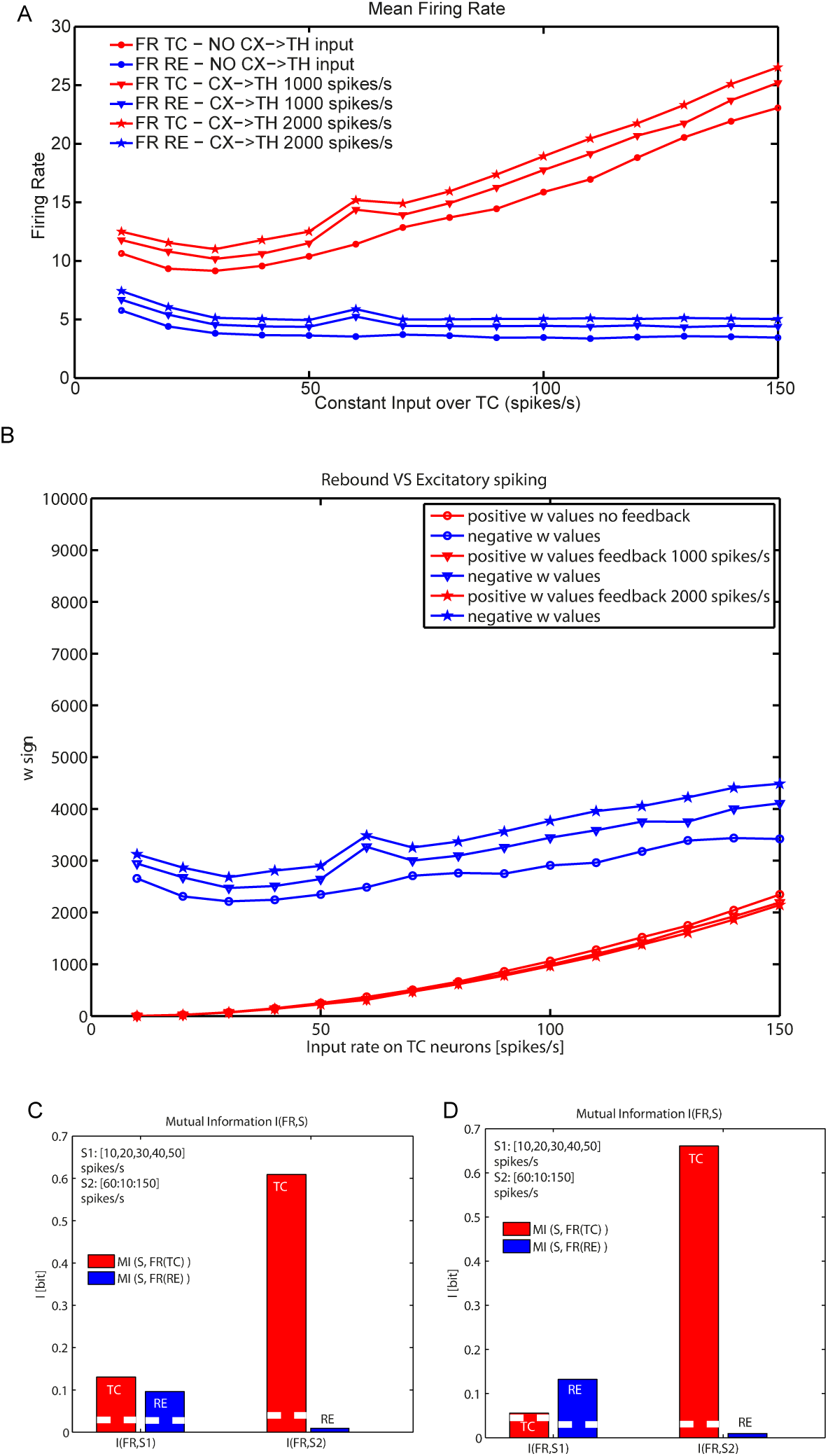
Bursting and tonic modes displayed by the TC-RE network with RE-RE clustering as a function of external input on TC neurons for different corticothalamic inputs. (A) Firing rate of TC (red) and RE (blue) neurons as a function of the external driving input impinging on TC neurons for different corticothalamic input amplitudes. (B) Number of positive (depolarizing, red) and negative (rebound, blue) *w* values of TC spikes for different corticothalamic inputs. The *w* values are averaged across 100 trials for each external stimulus. (C, D) Mutual Information carried by the firing rate of TC (red) and RE (blue) neurons with a cortico-thalamic input of (C) 1000 spikes/s and (D) 2000 spikes/s. I is calculated between the set of increasing sensory stimuli (10 – 150 spikes/s) and the neural response given by the firing rate. The white dashed lines in the bars refer to the significance threshold (*p* < 0.05, bootstrap test). Measures are averaged over 100 trials for each external stimulus. The synaptic strengths are respectively: *g_RE_*_→_*_TC_* = 300 *μ*S, *g_TC→RE_* = 200 *μ*S and *g_RE_*_→_*_RE_* = 300 *μ*S.

## Discussion

We have presented an adaptive exponential integrate-and-fire (aeIF) network model that is able to reproduce spindle oscillations and the transition between a stimulus-insensitive and a stimulus-sensitive state of the thalamus. Coherently to what was shown experimentally through direct optogenetic stimulation [18], in our model spindle oscillations are generated by RE activation leading to TC bursts as rebound from inhibition. Our simulations suggest that (i) these oscillations are stable for a specific range of RE-RE connection clustering, (ii) for external stimuli below a given threshold the network is in a purely rebound-bursting state insensitive to external stimuli, while when this threshold is crossed there is a non-zero contribution of the spikes due to depolarization, and this makes the TC neurons (and not the RE neurons) of the network sensitive to the stimulus intensity coherently with experimental observation.

### Advantages and limitations of aeIF models

Choosing a simple model for the single neurons allowed us to focus on capturing the network effects. This choice also opens a number of interesting perspectives: due to their relative simplicity, IF models can be tackled analytically [45, 46], and facilitate the search for basic canonical computations [47]. Finally, most primary sensory cortex network models are built on IF neurons [48, 49], and hence aeIF neurons seem a more coherent choice to build models of corticothalamic interactions [50]. In our model, the switch from inhibitory-rebound-driven activity to depolarization-driven firing is proposed to represent a switch from sleep to awake state [7]. The information analysis shown in Figs. 6–S4 shows the separation between a stimulus-independent state (sleep) and a stimulus-sensitive state (wakefulness). We did not directly deal with the role of thalamus, and in particular RE neuronal activity, in attention, to which a wealth of works have been devoted [9, 10] after the seminal intuition of Crick [2]. To compare our model results with these experimental observations we should (1) contrast different states inside the awake regime, and (2) take into account the temporal structure of the TC spike trains rather than their rate alone. This is certainly feasible on the ground of the results presented here, but is beyond the scope of this paper. We emphasize that our model is based on single-neuron models that are much simpler than those used previously. Although this has a number of advantages as discussed above, some features of thalamic behavior that are captured by more detailed models are not reproduced by our model. For instance, our spindle oscillations constitute a stable state, both in small and large TC-RE networks, and do not reproduce the wax-and-wane dynamics that has been observed experimentally [51], and which has been reproduced by more detailed models that take explicitly into account the dynamics of hyperpolarization-activated cation currents [52].

### TC-RE loop studies

A recent computational paper [25] investigated the role of TC-RE interactions from a perspective complementary to the one discussed in this paper, using a Hodgkin-Huxley model much more detailed than the aeIF adopted here, and limiting the investigation only to minimal loops such as those we described here (Fig. 2). Notwithstanding the higher realism of their model, the functional properties at the single-neuron level are similar to those described here (compare the two Figs. 1 of the two works). Moreover, Willis and colleagues highlighted the fact that open-loops between TC and RE neurons might play a functional role in the thalamus, and indeed in our full network (Fig. 4 and following) both open and closed TC-RE loops are taken into account. In a recent paper [21], Brown and collaborators stimulated optogenetically RE neurons, simultaneously recording from them. They found that the majority of those neurons (10/17) decreased significantly their firing rate, and only a minority of them (4/17) displayed a significant increase. At the same time they found that the activity of the TC neurons was inhibited, with functional consequences on the cortex. The interpretation of the authors was that a small increase in RE activity was sufficient to inhibit TC activity. Our model offers a simpler explanation: since most TC neurons fire due to hyperpolarization rebound, a decrease in RE activity can be associated to a decrease in TC firing (see Fig. 6). Indeed, stimulating RE neurons has been shown to alter the temporal structure of TC neuron firing, without changing their average firing rate [18].

### Perspectives

The present work has been focused on the thalamus, where we have taken into account only stable external inputs from periphery to TC neurons or from the cortex to RE neurons. Preliminary analysis suggested that an accurate description of thalamocortical inputs and corticothalamic feedbacks required a separate study. In the future this network will be integrated in a full corticothalamic model comprising a primary visual cortex network (inspired by previous works [53, 54]). The following step will be to take into account (i) the layered structure of the cortex [48] and (ii) areas of the thalamus and the cortex associated to different sensory receptive fields and their interactions. Another interesting continuation of this work would be to contribute to the open challenge of modeling the Local Field Potential of the thalamus [55]. We recently showed [56] that an integrate-and-fire model like the one presented here can be combined with morphological data and transmembrane current simulation [57] to capture the LFP dynamics in a patch of cortex. Since morphological data are available for the thalamus, a similar procedure can be applied to the network introduced here, and would hopefully shed light on the way extracellular signals and neural activity are linked in this area, thus enhancing the possibility of experimental validations of the thalamic models. The potential applications of this work include the study of the consequences of deep brain stimulation (DBS). Thalamic DBS has been shown to contribute to the symptom mitigation of a variety of neural diseases including Parkinson [58] and Tourette’s syndrome [59]. However, the precise mechanisms of this mitigation are not completely clear, nor is the procedure to design specific trains of stimulations suited for different patients/conditions. Neural models are already exploited to test DBS patterns [60]. We think that a simple yet efficient model like the one presented here can valuably contribute to this field.

## Materials and Methods

### Computational model

We have used the *adaptive exponential integrate-and-fire* (*aeIF*) model [27], which is an evolution of a two-variable integrate-and-fire (IF) model proposed by Izhikevich [31], and it is enriched by an exponential non-linearity around the spike threshold, as in the exponential IF model of Fourcaud-Trocme et al. [32]. The combination of these two models leads to the aeIF formulated by Brette and Gerstner [27].

### Single neuron model

According to the aeIF model, the equations describing the evolution of membrane voltage of neurons in the thalamus are:

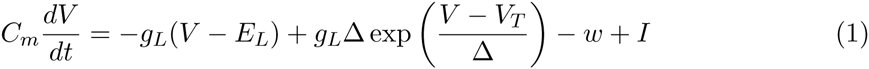

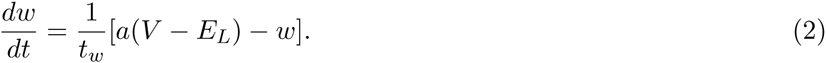

The first equation describes the evolution of the membrane voltage: the capacitive current through the membrane with capacitance *C_m_* = 1 nF equals the ionic currents, the adaptation current *w* and the input current *I*. The ionic currents are the ohmic leak current defined by the resting leak conductance *g_L_* = 0.05 *μ*S and the resting voltage potential *E_L_* = −60 mV, and the exponential term which reproduces the *Na^+^ −* current that is responsible for the generation of spikes. With this term we assume that the activation of *Na^+^ −*channels is instantaneous (thus neglecting their activation), with Δ denoting the steepness of the exponential approach to threshold, taken equal to Δ = 2.5 mV, and *V_T_* = −50 mV is the threshold potential. The membrane time constant is *τ_m_* = *C_m_/g_L_*. When *V* is pushed over the threshold, the exponential term provides a positive feedback and a spike is emitted (occurring ideally at the time when *V* diverges towards infinity), and the voltage is instantaneously reset to *V_r_* = −60 mV. After the spike, the neuron cannot spike again during a refractory period (2.5 ms).

The second equation describes the dynamics of the adaptation variable *w*, with time constant *τ_w_* = 600 ms. The parameter *a* (in *μ*S) quantifies a conductance that mediates subthreshold adaptation, while the increment *b* (in *n*A) at each spike takes into account spike-triggering adaptation (it regulates the strength of adaptation). When the input current *I* to the neuron at rest reaches a critical value (1000 mA) the resting state is destabilized, leading to repetitive spiking for large regions of parameter space [61]. Without adaptation (*a* = *b* = 0) the model produces tonic spiking. Neurons in general can show a reduction in the firing frequency of their spike response if they are stimulated with a square pulse or step, known as spike frequency adaptation (SFA). With this model, an increase of *a* or *b* leads to SFA, characterized by a gradual increase in the inter-spike interval (IS) until a steady-state spike frequency is reached. For *b* = 0, the model generates responses with a negative level of adaptation similar to the fast-spiking (FS) cells encountered in the cortex, often classified as inhibitory neurons. The strength of adaptation can be modulated by varying the parameter *b*, to get weakly adapting cells [61].

In order to reproduce the peculiar properties of TC and RE when operating in bursting mode, we adopted specific values of *a* and *b*. When they are in their tonic mode, TC and RE neurons behave similarly to excitatory regular spiking (RS) neurons and inhibitory fast spiking (FS) neurons found in the cortex. With *a* = 0.4 *μ*S, *b* = 0.02 *μ*A, neurons display bursting activity in response to both depolarizing and hyperpolarizing stimuli typical of RE neurons. In contrast, with *a* = 0.2 *μ*S, *b* = 0.0 *μ*A, neurons display responses with moderate adaptation and strong rebound bursts, like TC neurons. RE and TC neurons can display different regimes (beyond bursting and tonic, fast spiking (FS), regular spiking (RS)) by tuning the parameters *a* and *b* [31, 61, 62].

### Thalamic network model

The network is made of TC and RE cells, endowed with intrinsic properties and topographic connectivity specific to the thalamus [26]. Here we considered a network of 500 neurons, half of which are TC neurons and the other half being RE neurons. Given that thalamic interneurons do not contribute to the development of internal dynamics such as oscillations, they are neglected. Axonal projections within the thalamic circuitry are local but sparse. The excitatory projections from TC to RE had a connection probability of 1%, while RE to TC inhibitory projections were more dense, with a connection probability of 4%. The same density was assumed from inhibitory connections between RE cells. The structural connectivity is built according to *small world algorithm* of Strogatz and Watts [41] in which neurons are first built in a ring network and then randomly rewired with rewiring probability RP. In fig. 4 we introduced different degree of clustering (by tuning the rewiring probability (RP)) and according to the results we adopted *RP* = 0.25 for the continuation of the analysis. The network model was constructed based on this aeIF model, according to the following equations [26]:

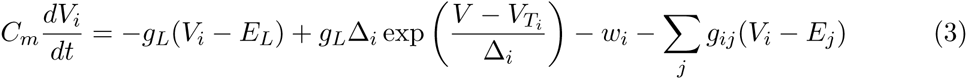

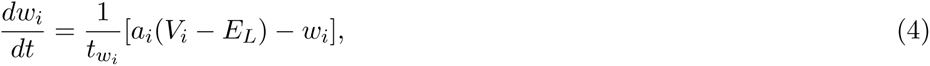

where *V_i_* is the membrane potential of neuron i, and all parameters are as in Eqs. (1)-(2), but were indexed to allow variations according to the cell type. The term 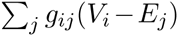 accounts for the synaptic current coming from the neighboring neurons impinging on a neuronal cell, where *g_ij_* is the conductance of the synapse from neuron *j* to neuron *i* (which can be zero), and *E_j_* is the reversal potential of the synapse (*E_j_* = 0 mV for excitatory synapses and <80 mV for inhibitory synapses). Synaptic conductances are described by:

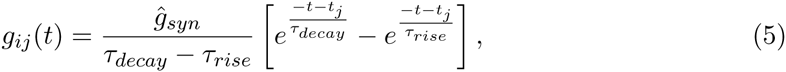

where *τ_decay_* and *τ_rise_* are the decay and rise synaptic time, respectively, and *ĝ_syn_* is constant and depends on the type of synapses and network (see table 1). Once the presynaptic cell fires, *g_ij_* exponentially increases up to a certain value, after which *g_ij_* decays exponentially with a fixed time constant (5 ms for excitation and 10 ms for inhibition). Different synaptic strengths are considered (see table 2), depending on the network type. If different values are considered, they are indicated in the captions of each figure. Synaptic delays are equal to 1 ms.

**Table 1.** Values of temporal rise and decay constants for RE and TC.

| | AMPA $\tau_{rise}$ | AMPA $\tau_{decay}$ | GABA $\tau_{rise}$ | GABA $\tau_{decay}$ |
| --- | --- | --- | --- | --- |
| Network 2 neurons | 0.4 ms | 5 ms | 0.4 ms | 20 ms |
| Network 4 neurons | 0.4 ms | 5 ms | 0.4 ms | $\mu = 20$ ms, $\sigma = 5$ ms |
| Network 500 neurons | 0.4 ms | 5 ms | 0.4 ms | 10 ms |

**Table 2.** Values of synaptic strengths for a network of 500 neurons.

| | $g_{RE \rightarrow TC}$ | $g_{TC \rightarrow RE}$ | $g_{RE \rightarrow RE}$ | $g_{ext \rightarrow TC}$ | $g_{CX \rightarrow RE}$ |
| --- | --- | --- | --- | --- | --- |
| Network 2 neurons | 200 – 800 μS | 10 – 60 μS | 200 – 800 μS | 1 μS | 1 μS |
| Network 4 neurons | 550 μS | 32 μS | 20 μS | 1 μS | 1 μS |
| Network 500 neurons | 300 μS | 200 μS | 300 μS | 5 μS | 1 μS |

To initiate activity, during the first 50 ms a number of randomly-chosen neurons were stimulated by an incoming current (with synaptic strength g= 40 *μ*S), representing an heterogenous Poisson train of excitatory presynaptic potential with an instantaneous event rate λ(*t*) that varies following an Ornstein-Uhlenbeck process:

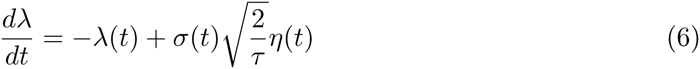

where *σ*(*t*) is the standard deviation of the noise and is set to 0.6 spikes/s. *τ* is set to 16 ms, leading to a power spectrum for the *λ* time series that is approximately flat up to a cut-off frequency 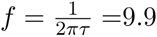 Hz. *η*(*t*) is a Gaussian white noise of mean zero and intensity unity. In simulations in which we did not take into account external input after 50 ms, no input was given to the network, and thus the activity states described here are self-sustained with no external input or added noise. The only source of noise was the random connectivity. In simulations in which we took into account external sensory inputs, after 5 s of self-sustained activity we injected for 10 s homogeneous Poisson processes with rate comprised between 10 and 150 spikes/s.

### Spectral analysis

We computed the power spectral density of LFPs and MUAs using the Welch method: the signal is split up into 32768 point segments with 50% overlap. The overlapping segments are windowed with a Hamming window. The modified periodogram is calculated by computing the discrete Fourier Transform, and then calculating the square magnitude of the result. The modified periodograms are then averaged to obtain the PSD estimate, which reduces the variance of the individual power measurements. Spectral quantities and phase coherence are averaged over 50 trials.

### Phase coherence

Phase coherence is calculated as in [63]:

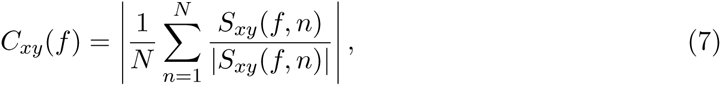

where *x* and *y* denote the two signals, and *S_xy_*(*f, n*) is the cross-spectrum between them. Since in each trial the cross spectral density is normalized by its amplitude, each term of the sum is a unit-length vector representation of the phase relation Δ*ϕ*(*f, n*). In other words, Δ*ϕ*(*f, n*) = *ϕ_y_ – ϕ_x_* is the phase lag between the two signals at frequency *f* in the data segment *n*. Hence *C_xy_*(*f*) quantifies how broad is the distribution of Δ*ϕ*(*f, n*) within the 2*π*-cycle. Averaging Δ*ϕ*(*f, n*) across all *N* data segments provides a mean angle Δ*ϕ*(*f*).

### Mutual information

We calculate the Mutual Information *I*(*S*; *FR*) between the set of stimuli *S* given by the external Poisson inputs with different rates described above and the response *FR*, firing rate, as follows. Given that we are interested in how the specific neurons encode and carry information, in this case we select as response the average firing rate *FR* over the whole stimulation (other responses such as the power spectrum can be considered [64]). We consider as stimuli different inputs with increasing amplitude (from 0 to 150 spikes/s) impinging on TC neurons. We compute the information between the stimulus *S* and the response firing rate as:

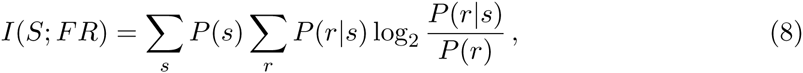

where *P*(*s*) is the probability of having a stimulus *s* (equal to the inverse of the total number of different external firing rates, which act as stimuli), *P*(*r*) is the probability of observing a firing rate *r* across all trials in response to any stimulus, and *P*(*r*|*s*) is the probability of observing a firing rate *r* in response to a single stimulus s. *I*(*S*; *FR*) quantifies the reduction of uncertainty about the stimulus that can be gained from observing a single-trial neural response, measured in units of bits (1 bit means a reduction of uncertainty of a factor of two) [43]. This measure allows us to evaluate how well the firing rate *r* of both type of neurons encodes the stimulus *s*.

An important issue to be solved regarding the calculation of the mutual information is that it requires knowledge of the full stimulus-response probability distributions, and obviously these probabilities are calculated from a finite number of stimulus-response trials. This leads to the so-called limited sampling bias, which constitutes a systematic error in the estimate of information. We used the method described in [65] to estimate the bias of the information quantity and then we checked for the residual bias by applying a *bootstrap procedure*, in which mutual information is calculated when the stimuli and responses are paired at random. If the information quantity is not zero (as it should be in the case of non-finite samples), this is an indication of the bias, and the bootstrap estimate of this error should be removed from the mutual information. After applying these procedures, the information quantity estimation could be defined as significant. Several toolboxes provide different bias-correction techniques, which allow accurate estimates of information theoretic quantities from realistically collectable amounts of data [66, 67]. In order to accomplish those tasks, we used the Information Breakdown Toolbox (ibTB), a MATLAB toolbox implementing several information estimates and bias corrections [67].

## Acknowledgments

This work was supported by the European Commission under ITN project NETT (FP7 contract 289146), by the Spanish Ministry of Economy and Competitiveness and FEDER (project FIS2012–37655-C02–01), and by the Generalitat de Catalunya (project 2014SGR0947). A. M. was supported by the NEBIAS European project (EUFP7-ICT-611687), by the PRIN/ HandBot Italian project (CUP: B81J12002680008; prot.: 20102YF2RY), and by the Italian Ministry of Foreign Affairs and International Cooperation, Directorate General for Country Promotion (Economy, Culture and Science) Unit for Scientific and Technological Cooperation, via the Italy-Sweden bilateral research project on “Brain network mechanisms for integration of natural tactile input patterns”. J.G.O. also acknowledges support from the ICREA Academia programme and from the “Maria de Maeztu” Programme for Units of Excellence in R&D (Spanish Ministry of Economy and Competitiveness, MDM-2014–0370).

## Supplementary Figures

**Figure S1.**
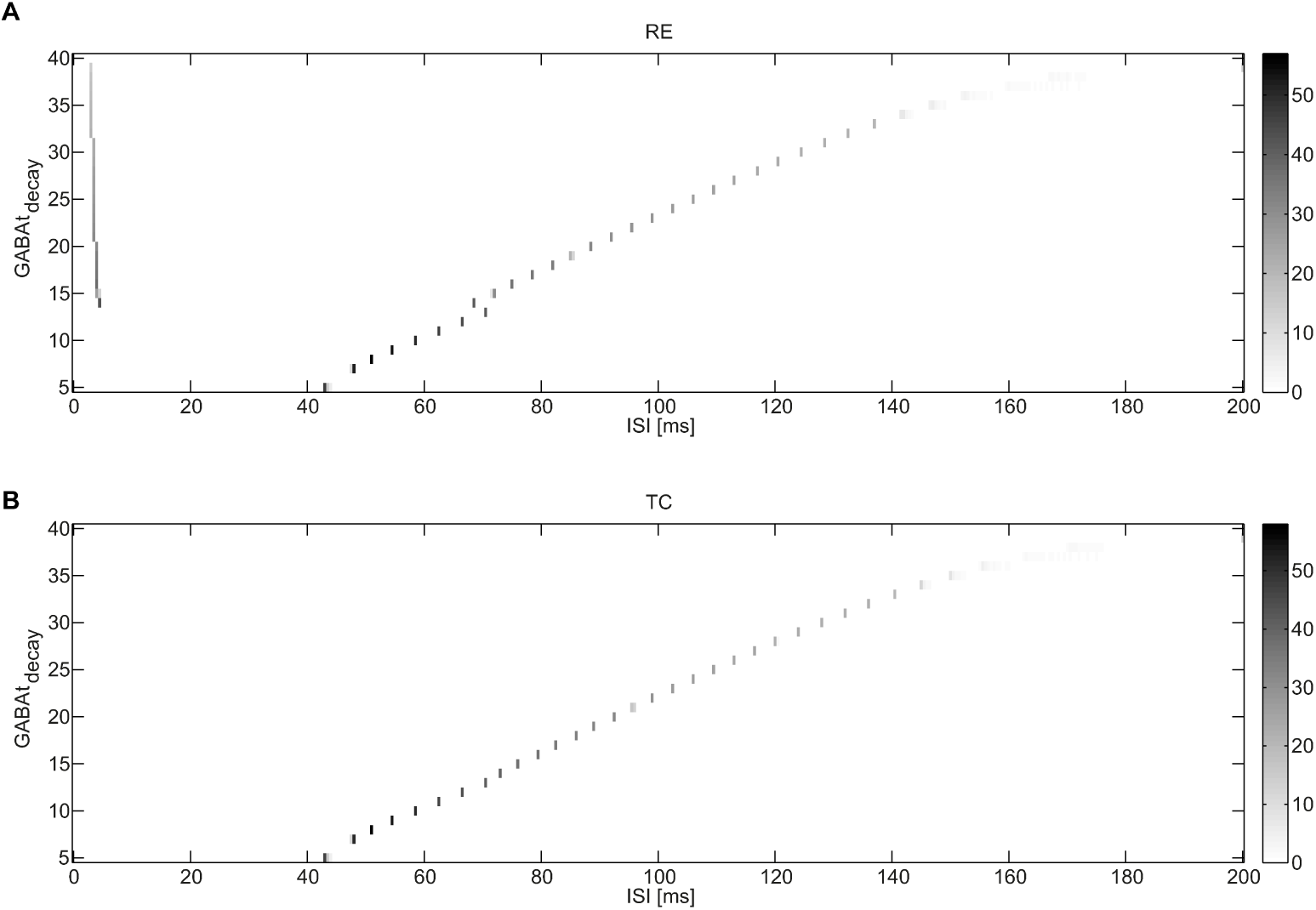
Effect of the GABA decay time on the two-neuron TC-RE loop. Interspike Interval (ISI) distribution of a TC-RE loop as a function of the GABA decay time *τ_decay_* for RE (A) and TC (B) neurons. *τ_decay_* varies between 5 and 40 ms. The synaptic strengths are respectively: *g_RE_*_→_*_TC_* = 550 *μ*S, *g_TC→RE_* = 32 *μ*S.

**Figure S2.**
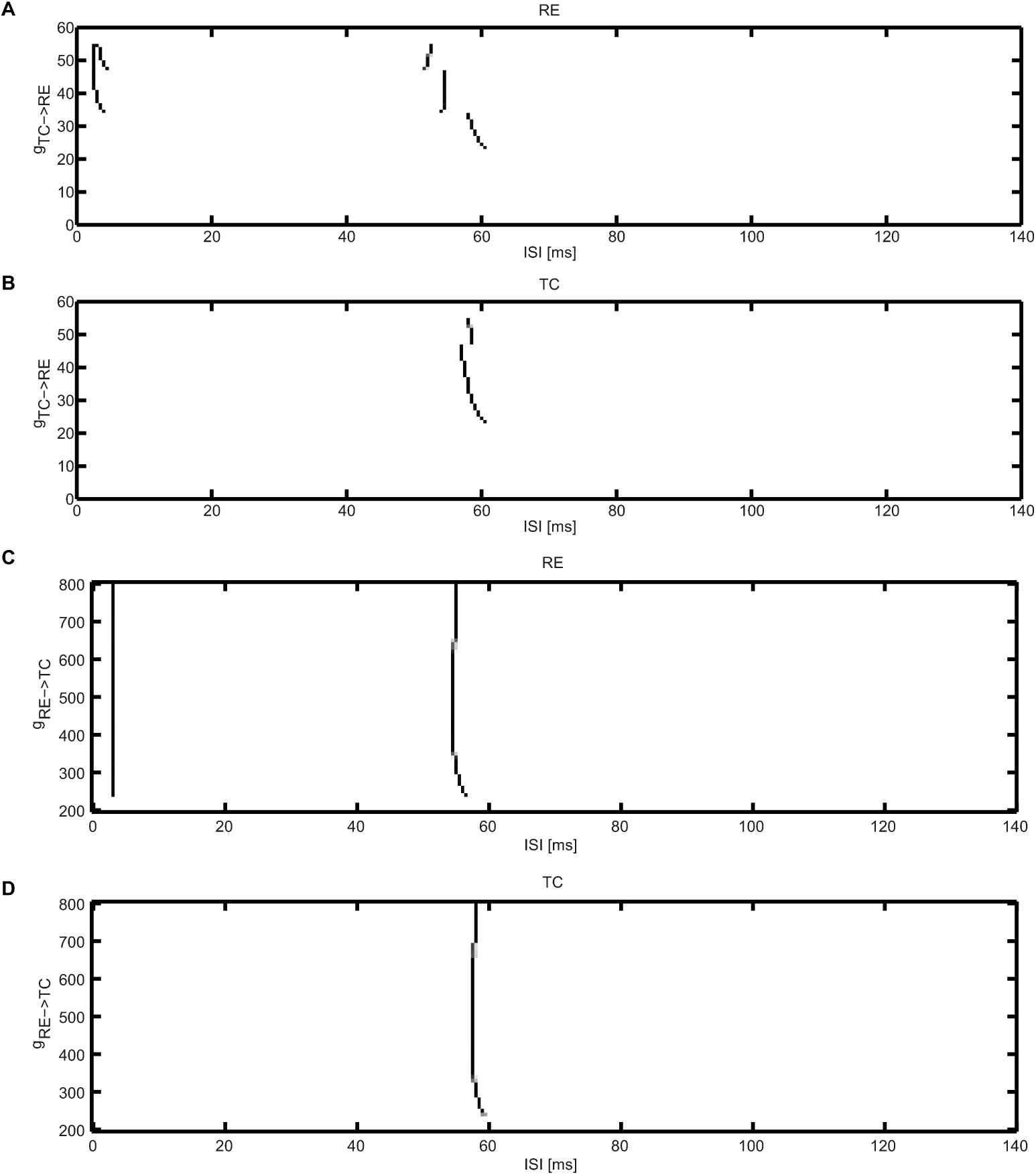
Effect of the synaptic strength on the two-neuron TC-RE loop. Interspike Interval (ISI) distribution of a TC-RE loop as a function of the synaptic strength for RE (A, C) and TC (B, D) neurons. As in fig. 2C, in A and B the value of *g_RE_*_→_*_TC_* is appropriately set to 550 *μ*S in order to support self-sustained activity, while *g_TC→RE_* varies between 10 *μ*S and 60 *μ*S. In C and D, the value of *g_TC→RE_* is chosen equal to 40 *μ*S to reproduce the two-spike bursting dynamical regime, while *g_RE_*_→_*_TC_* varies between 200 *μ*S and 800 *μ*S. GABA decay *τ_decay_* is set equal to 10 ms in the two cases.

**Figure S3.**
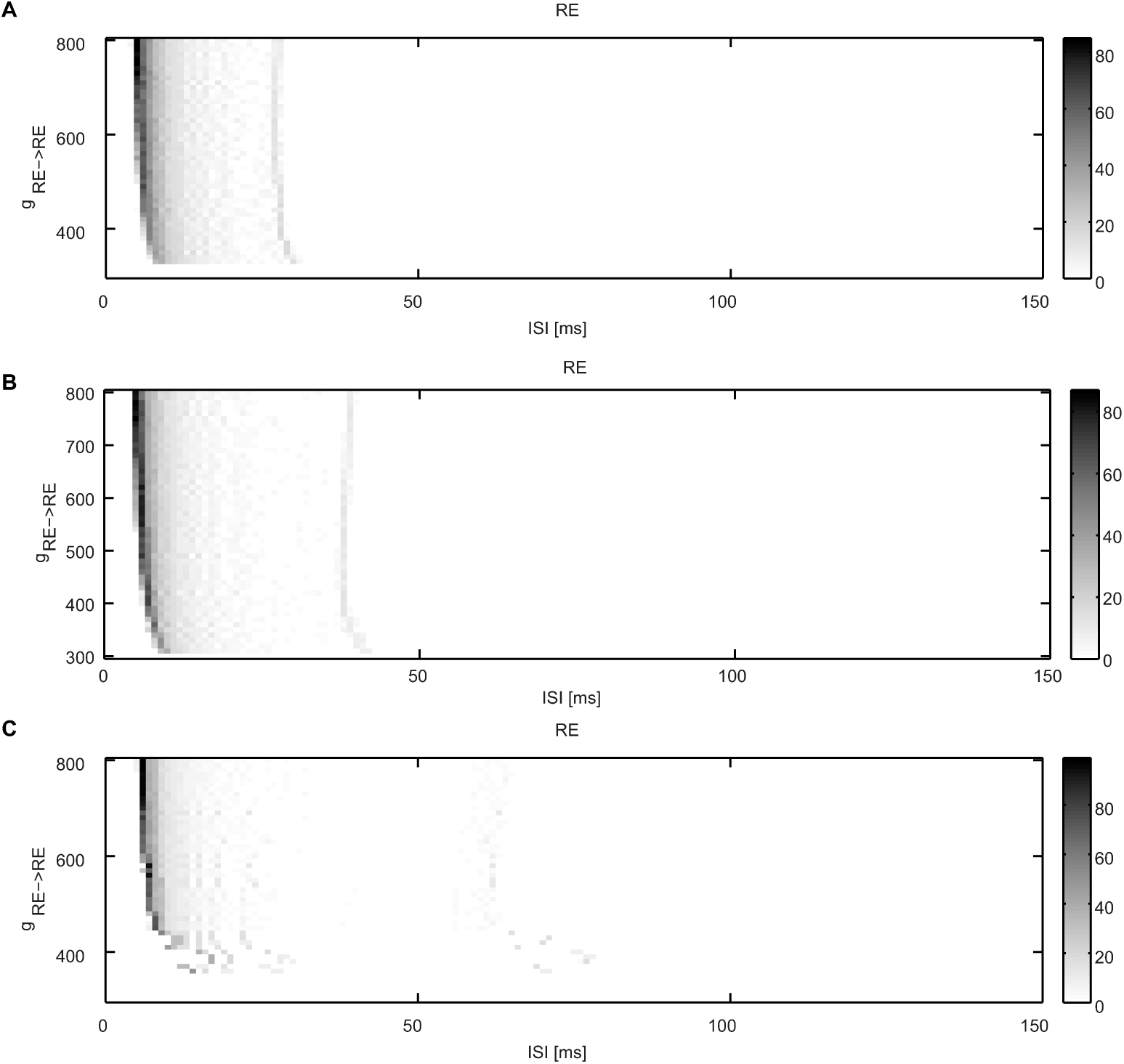
Effect of the synaptic strengths on the two-neuron RE-RE motif. Interspike interval (ISI) distribution of a minimal purely reticular RE-RE motif as a function of the synaptic strength *g_RE_*_→_*_RE_* for three different values of the GABA decay time: (A) 5 ms, (B) 10 ms, (C) 20 ms. *g_RE_*_→_*_TC_* varies between 300 *μ*S and 800 *μ*S.

**Figure S4.**
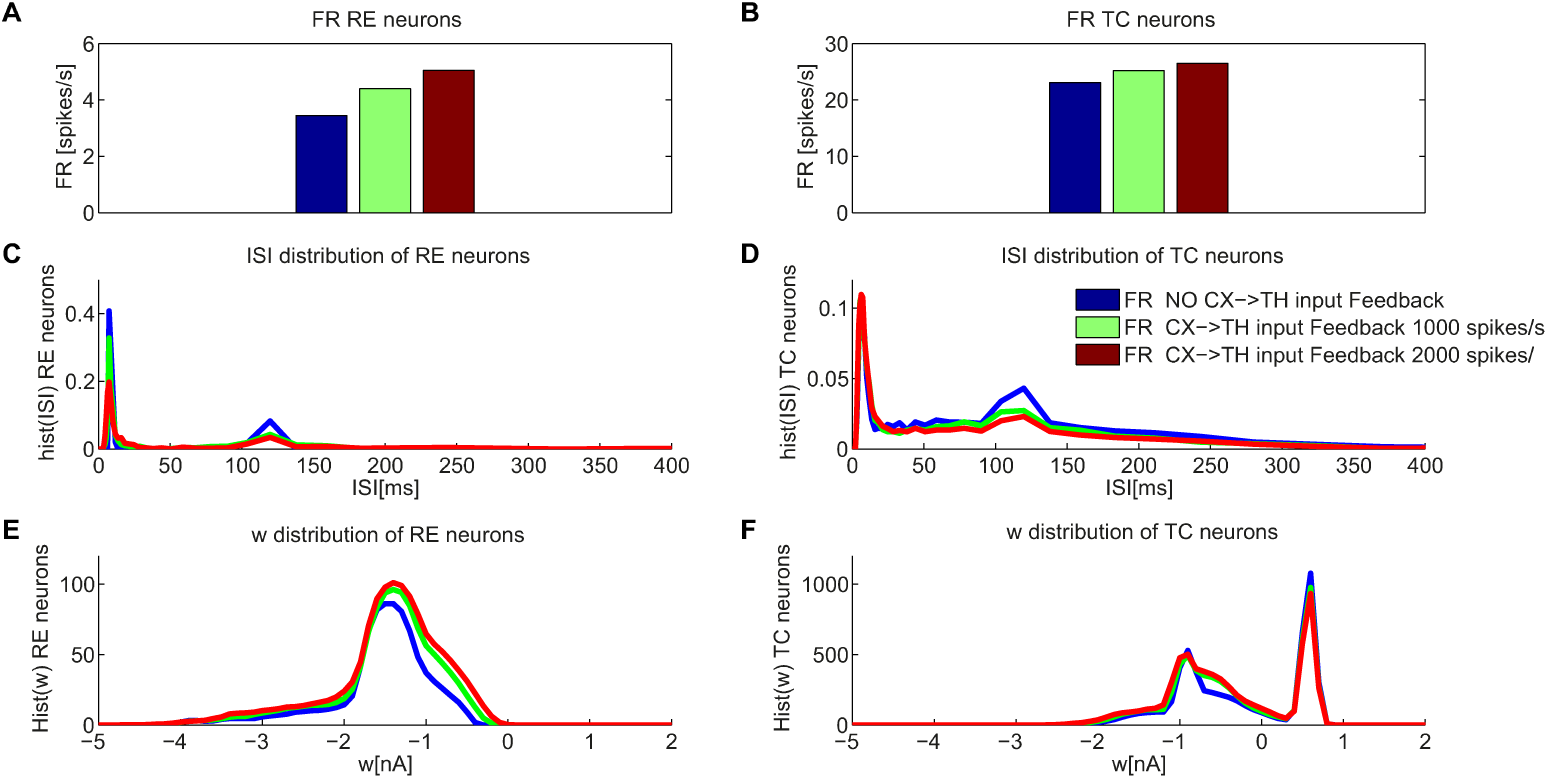
Influence of corticothalamic input on a full TC-RE network. (A) Firing rate, (B) ISI distribution, and (C) distribution of the adaptation variable *w* of RE and TC neurons as a function of corticothalamic input. The external sensory input it set to 150 spikes/s. The synaptic strengths are respectively: *g_RE_*_→_*_TC_* = 300 *μ*S, *g_TC→RE_* = 200 *μ*S and *g_RE_*_→_*_RE_* = 300 *μ*S. Bars colors in panels (A) and (B) coincide with the lines colors in the other panels.

